# Identifying genetic variants that affect viability in large cohorts

**DOI:** 10.1101/085969

**Authors:** Hakhamanesh Mostafavi, Tomaz Berisa, Felix R Day, John R B Perry, Molly Przeworski, Joseph K Pickrell

**Affiliations:** Department of Biological Sciences, Columbia University, New York, NY, USA; New York Genome Center, New York, NY, USA; MRC Epidemiology Unit, Institute of Metabolic Science, University of Cambridge, Cambridge, UK; Department of Systems Biology, Columbia University, New York, NY, USA

## Abstract

A number of open questions in human evolutionary genetics would become tractable if we were able to directly measure evolutionary fitness. As a step towards this goal, we developed a method to examine whether individual genetic variants, or sets of genetic variants, currently influence viability. The approach consists in testing whether the frequency of an allele varies across ages, accounting for variation in ancestry. We applied it to the Genetic Epidemiology Research on Aging (GERA) cohort and to the parents of participants in the UK Biobank. Across the genome, we find only a few common variants with large effects on age-specific mortality: tagging the *APOE* ε4 allele and near *CHRNA3*. These results suggest that when large, even late onset effects are kept at low frequency by purifying selection. Testing viability effects of sets of genetic variants that jointly influence one of 42 traits, we detect a number of strong signals. In participants of the UK Biobank study of British ancestry, we find that variants that delay puberty timing are enriched in longer-lived parents (*P*~6×10^−6^ for fathers and *P*~2×10^−3^ for mothers), consistent with epidemiological studies. Similarly, in mothers, variants associated with later age at first birth are associated with a longer lifespan (*P*~1×10^−3^). Signals are also observed for variants influencing cholesterol levels, risk of coronary artery disease, body mass index, as well as risk of asthma. These signals exhibit consistent effects in the GERA cohort and among participants of the UK Biobank of non-British ancestry. Moreover, we see marked differences between males and females, most notably at the *CHRNA3* locus, and variants associated with risk of coronary artery disease and cholesterol levels. Beyond our findings, the analysis serves as a proof of principle for how upcoming biomedical datasets can be used to learn about selection effects in contemporary humans.

## Introduction

A number of central questions in evolutionary genetics remain open, in particular for humans. Which types of variants affect fitness? Which components of fitness do they affect? What is the relative importance of directional and balancing selection in shaping genetic variation? Part of the difficulty is that our understanding of selection pressures acting on the human genome is based either on experiments in fairly distantly related species or cell lines or on indirect statistical inferences from patterns of genetic variation [1-3].

The statistical inferences rely on patterns of genetic variation in present day samples (or very recently, in ancient samples [4]) to identify regions of the genome that appear to carry the footprint of positive selection [2]. For example, a commonly used class of methods asks whether rates of non-synonymous substitutions between humans and other species are higher than expected from putatively neutral sites, in order to detect recurrent changes to the same protein [5]. Another class instead relies on polymorphism data and looks for various footprints of adaptation involving single changes of large effect [6]. These approaches detect adaptation over different timescales and, likely as a result, suggest quite distinct pictures of human adaptation [1]. For example, approaches that are sensitive to selective pressures acting over millions of years have identified individual chemosensory and immune-related genes (e.g., [7]). In contrast, approaches that are most sensitive to selective pressures active over thousands or tens of thousands of years have revealed strong selective pressures on individual genes that influence human pigmentation (e.g., [8-10]), diet [11-13], as well as sets of variants that shape height [14-16]. Even more recent still, studies of contemporary populations have suggested that natural selection has influenced life history traits like age at first childbirth as well as educational attainment over the course of the last century [17-23].

Because these approaches are designed (either explicitly or implicitly) to be sensitive to a particular mode of adaptation, they provide a partial and potentially biased picture of what variants in the genome are under selection. In particular, most have much higher power to adaptations that involve strongly beneficial alleles that were rare in the population when first favored and will tend to miss selection on standing variation or adaptation involving many loci with small beneficial effects (e.g., [24-27]). Moreover, even where these methods identify a beneficial allele, they are not informative about the components of fitness that are affected or about possible fitness trade-offs between sexes or across ages.

In line with Lewontin’s proposal to track age-specific mortality and fertility of hundreds of thousands of individuals for the study of natural selection [28], we introduce a more direct and, in principle, comprehensive way to study adaptation in humans, focusing on *current* viability selection. Similar to the approach that Allison took in comparing frequencies of the sickle cell allele in newborn and adults living in malarial environments [29], we aim to directly observe the effects of genotypes on survival, by taking advantage of the recent availability of genotypes from a large cohort of individuals of different ages. Specifically, we test for differences in the frequency of an allele across individuals of different ages, controlling for changes in ancestry and possible batch effects. This approach resembles a genome-wide association study for longevity, yet does not focus on an endpoint (e.g. survival to an old age) but on any shift in allele frequencies with age. Thus, it allows the identification of possible non-monotonic effects at different ages or sex differences. Any genetic variant that affects survival by definition has a fitness cost, even if the cost is too small to be effectively selected against (depending on the effective population size, the age structure of the population and the age at which the variant exerts its effects [30]). Of course, a genetic variant can influence fitness without influencing survival, through effects on reproduction or inclusive fitness. Our approach is therefore considering only one of the components of fitness that are likely important for human adaptation.

As a proof of principle, we applied our approach to two recent datasets: to 57,696 individuals of European ancestry from the Resource for Genetic Epidemiology Research on Aging (GERA) Cohort [31, 32] and, by proxy [33-35], to the parents of 117,649 individuals of British ancestry surveyed as part of the UK Biobank [36]. We did so for individual genetic variants, then jointly for sets of variants previously found to influence one of 42 polygenic traits [37-40].

## Results

### A method for testing for differences in allele frequencies across age bins

If a genetic variant does not influence viability, its frequency should be the same in individuals of all ages. We therefore test for changes in allele frequency across individuals of different ages, while accounting for systematic differences in the ancestry of individuals of different ages (for example, due to migration patterns over decades) and genotyping batch effects. We use a logistic regression model in which we regress each individual’s genotype on their age bin, their ancestry as determined by principal component analysis (PCA) (Figure S1), and the batch in which they were genotyped (see Materials and Methods for details). In this model, we treat age bin as a categorical variable; this allows us to test for a relationship between age and the frequency of an allele regardless of the functional form of this relationship. We also test a model with an interaction between age and sex, to assess whether a variant affects survival differently in the two sexes.

We first evaluated the power of this method using simulations. We considered three possible trends in allele frequency with age: (i) a constant frequency up to a given age followed by a steady decrease, i.e., a variant that affects survival after a given age (e.g., variants contributing to late-onset disorders), (ii) a steady decrease across all ages for a variant with detrimental effect throughout life, and (iii) a U-shape pattern in which the allele frequency decreases to a given age but then increases, reflecting trade-offs in the effects at young and old ages, as hypothesized by the antagonistic pleiotropy theory of aging [41] or as may be seen if there are protective alleles that buffer the effect of risk alleles late in life [42] (Figure 1). In all simulations, we used sample sizes and age distributions that matched the GERA cohort (Figure S2). For simplicity, we also assumed no population structure or batch effects across age bins (Materials and Methods). For all trends, we set a maximum of 20% change in the allele frequency from the value in the first age bin (Figure 1).

**Figure 1.**
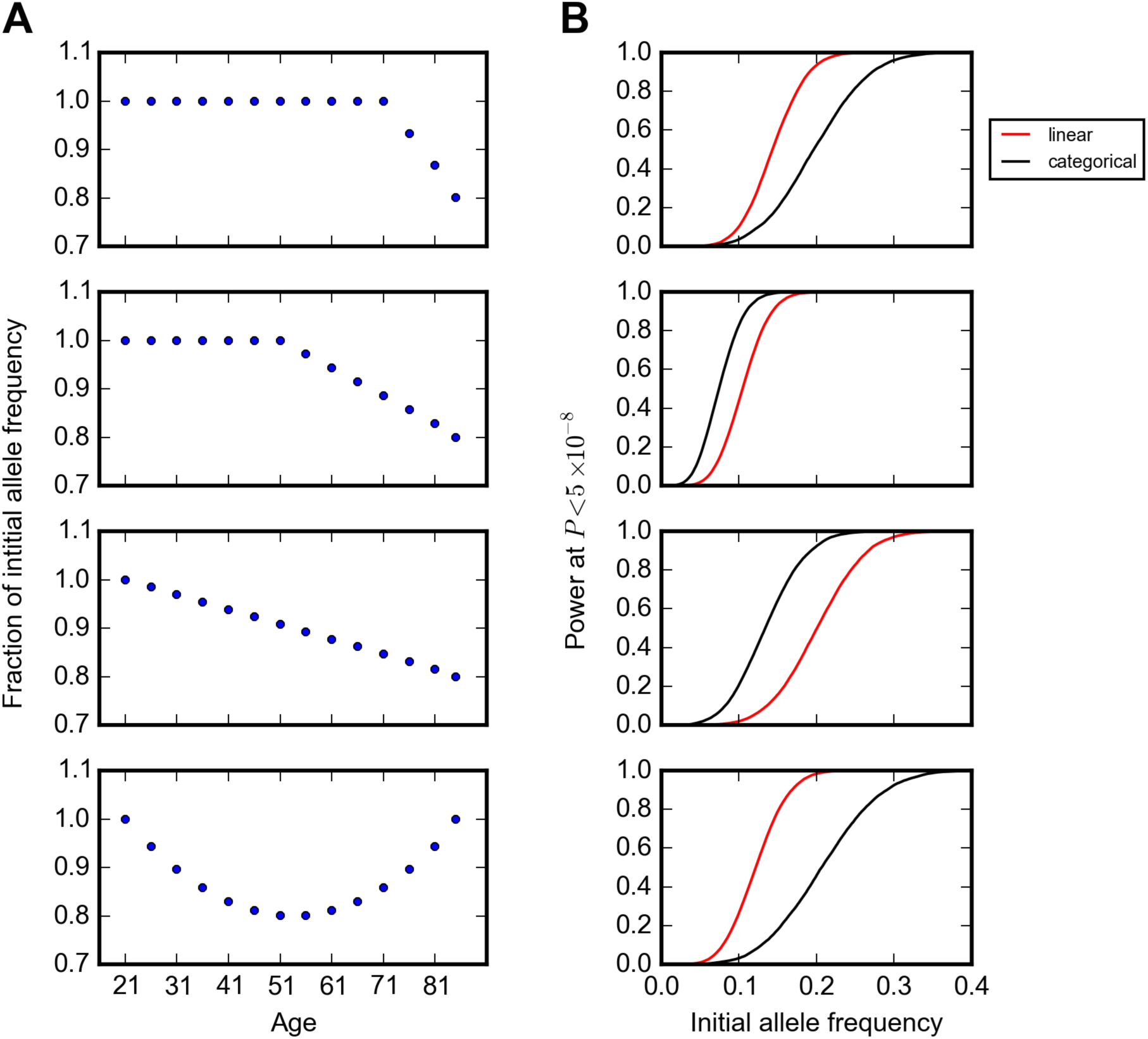
Power of the model to detect changes in allele frequency with age. (A) Trends in allele frequency with age considered in simulations. The y-axis indicates allele frequency normalized to the frequency in the first age bin. (B) Power to detect the trends in (A) at *P* < 5×10^−8^, given the sample size per age bin in the GERA cohort (Figure S2 and total sample size of 57,696). Shown are results using models with age treated as a categorical (black) or an ordinal (red) variable, assuming no change in population structure and batch effects across age bins. The curves show simulation results sweeping allele frequency values with an increment value of 0.001 (1000 simulations for each allele frequency) smoothed using a Savitzky-Golay filter using the SciPy package [80].

Because of the age distribution of individuals in the GERA cohort (Figure S2), our power to detect the trend is greater when most of the change in allele frequency occurs at middle age (Figure 1). For example, for an allele with an initial allele frequency of 15% that begins to decrease in frequency among individuals at age 20, age 50, or age 70 years, there is around 20%, 90% and 60% power, respectively, to detect the trend at *P* < 5×10^−8^, the commonly-used criterion for genome-wide significance [43]. We also experimented with a version of the model where the age bin is treated as an ordinal variable; as expected, this model is more powerful if there is a linear relationship between age and allele frequency (Materials and Methods). Since in most cases, we do not know the functional form of the relationship between age and allele frequency *a priori*, we used the categorical model for all analyses, unless otherwise noted.

In the UK Biobank, all individuals were 45-69 years old at enrollment, so the age range of the participants is restricted and our method has low power. However, the UK Biobank participants reported the survival status of their parents: age of the parents if alive, or age at which their parents died; following recent studies [33-35], we therefore used these values (when reported) instead in our model. In this situation, we are testing for correlations between an allele frequency and father’s or mother’s age (if alive) or age at death (if deceased). This approach obviously comes with the caveat that children inherit only 50% of their genome from each parent and so power is reduced (e.g., [44]). Furthermore, the patterns expected when considering individuals who have died differ subtly from those generated among surviving individuals. Notably, when an allele begins to decline in frequency starting at a given age (Figure 1A), there should be an *increase* in the allele frequency among individuals who died at that age, followed by a decline in frequency, rather than the steady decrease expected among surviving individuals (Figure S3, see Materials and Methods for details). In a first analysis, we therefore focused on the majority of participants that reported father’s or mother’s age at death, 88,595 and 71,783 individuals, respectively. We compared the results of this approach with the results of a Cox proportional hazard model [45], which allows us to include individuals who reported their parents to be alive, but has the disadvantage of assuming fixed effects across all ages.

We further adapted this model to allow us to test for changes in frequency at sets of genetic variants jointly. Many phenotypes of interest, from complex disease risk to anthropomorphic and life history traits such as age at menarche, are polygenic [46, 47]. If a polygenic trait has an effect on fitness, either directly or indirectly (i.e., through pleiotropic effects), the individual loci that influence the trait may be too subtle in their survival effects to be detectable with current sample sizes. We therefore investigate whether there is a shift across ages in *sets* of genetic variants that were identified as influencing a trait in genome-wide association studies (GWAS) (Table S1). Specifically, for a given trait, we calculate a polygenic score for each individual based on trait effect sizes of single variants previously estimated in GWAS and then test whether the scores vary significantly across 5-year age bins (see Materials and Methods for details). These scores are calculated under an additive model, which appears to provide a good fit to GWAS data [48].

If a polygenic trait is under stabilizing selection (e.g., human birth weight [49]), i.e., an intermediate polygenic score is optimal, no change in the mean value of polygenic score across different ages is expected. However, if extreme values of a trait are associated with lower chance of survival, the spread of the polygenic scores should decrease with age. To consider this possibility, we tested whether the squared difference of the polygenic scores from the population mean is significantly associated with survival (see Materials and Methods for details).

### Testing for changes in allele frequency at individual genetic variants

We first applied the method to the GERA cohort, using 9,010,280 filtered genotyped and imputed autosomal biallelic single-nucleotide polymorphisms (SNPs) and indels. We focused on a subset of filtered 57,696 individuals who we confirmed to be of European ancestry by PCA (see Materials and Methods, Figures S4 and S5). The ages of these individuals were reported in bins of 5 year intervals (distribution shown in Figure S2). We tested for significant changes in allele frequencies across these bins. For each variant, we obtained a *P* value comparing a model in which the allele frequency changes with age to a null model. No inflation was observed in the quantile-quantile plot (Figure S6A), indicating that, for common variants at least, our control for population structure (and other potential confounders) is sufficient. To illustrate this point, we looked at the lactose intolerance linked SNP rs4988235 within the *LCT* locus, which is among the most differentiated variants across European populations [11]; the trend in the expected allele frequency based on the null model (i.e., accounting for confounding batch effects and changes in ancestry) tracks the observed trend quite well (Figure S7).

By our approach, all variants that reached genome-wide significance (*P* < 5×10^−8^) reside on chromosome 19 near the *APOE* gene (Figure 2A and Figure S8). This locus has previously been associated with longevity in multiple studies [50, 51]. The ε4 allele of the *APOE* gene is known to increase the risk of late-onset Alzheimer’s disease (AD) as well as of cardiovascular diseases [52, 53]. We observe a monotonic decrease in the frequency of the T allele of the ε4 tag SNP rs6857 (C, protective allele; T, risk allele) beyond the age of 70 years old (Figure 2B). This trend is observed for both the heterozygous and homozygous risk variants (Figure S9), and for both males and females (Figure S10). No variant reached genome-wide significance testing for age by sex interactions (quantile-quantile plot shown in Figure S6B).

**Figure 2.**
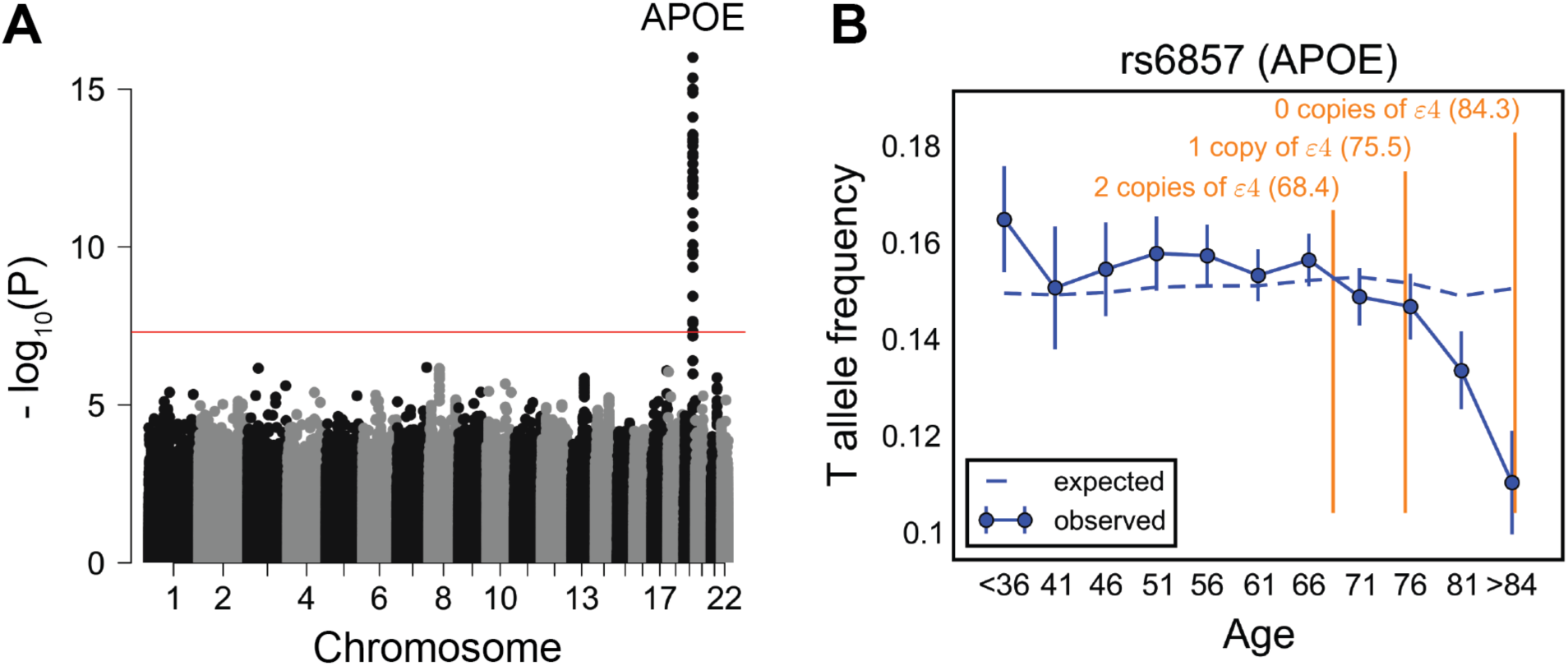
Testing for the influence of single genetic variants on age-specific mortality in the GERA cohort. (A) Manhattan plot of *P* values for the change in allele frequency with age. Red line marks the *P* = 5×10^−8^ threshold. (B) Allele frequency trajectory of rs6857, a tag SNP for *APOE* ε4 allele, with age. Data points are mean frequencies of the risk allele within 5-year interval age bins (and 95% confidence interval), with the center of the bin indicated on the x-axis. Bins with ages below 36 years are merged into one bin because of the relatively small sample sizes per bin. The dashed line shows the expected frequency based on the null model accounting for confounding batch effects and changes in ancestry (see Materials and Methods). In orange are the mean ages of onset of Alzheimer’s disease for carriers of 0, 1 or 2 copies of the *APOE* ε4 allele [52].

We further investigated the trends in frequency with age for the other two major *APOE* alleles defined by rs7412 and rs429358 SNPs: ε2 (rs7412-T, rs429358-T) and ε3 (rs7412-C, rs429358-T), while ε4 is (rs7412-C, rs429358-C) [54]. Unlike the ε4 allele, ε2 carriers are suggested to be at lower risk of Alzheimer’s disease, cardiovascular disease, and mortality relative to the ε3 carriers [50, 54]. We focused on a subset of 38,703 individuals with unambiguous counts of each *APOE* allele. There is a significant change in the frequency of the ε4 allele with age in this subset (*P*~6×10^−12^), similar to the trend observed for the tag SNP rs6857 (Figure S11). The ε3 allele shows the reverse trend, with a significant, monotonic increase in frequency beyond age of 70 years old (*P*~2×10^−8^) (Figure S11). The enrichment of the ε3 allele in elderly individuals can be explained by the corresponding depletion of the ε4 allele, however, so does not necessarily imply an independent, protective effect of ε3. The frequency of the ε2 allele does not change significantly with age (*P*~0.2), possibly reflecting low power, given its allele frequency of ~0.06 (Figure S11).

We considered the possibility that some unobserved confounding variable was driving the strength of this signal at *APOE*. Since there are two genotyped SNPs with signals similar to rs6857 within the locus, genotyping error seems unlikely to be driving the pattern (Figure S8). Another concern might be a form of ascertainment bias, in which individuals with Alzheimer’s disease are underrepresented in the Kaiser Permanente Medical Care Plan. However, there is no correlation in these data between the amount of time that an individual has been enrolled in this insurance plan and the individual’s *APOE* genotype (Figure S12). These observations, along with previously reported associations at this locus, argue that the allele frequency trends in Figure 2B are driven by effects of *APOE* genotype on mortality (or severe disability). Moreover, the effects that we identified are concordant with epidemiological data on the peak age of onset of Alzheimer’s disease given 0 to 2 copies of *APOE* ε4 [52]. This case not only serves as a positive control for our approach, it illustrates the resolution that it provides about age effects of genetic variants.

We estimated that we have ~93% power to detect the trend in allele frequency with age as observed for rs6857 (at a genome-wide significance level; see Materials and Methods). Using both versions of the model treating age bin as a categorical or an ordinal variable, we have similar power to detect other potential trends considered in Figure 1, for variants as common as rs6857 and with similar magnitude of effect on survival. Yet across the genome, only *APOE* variants show a significant change in allele frequency with age for both versions of the model (Figure 2 and Figure S13). Thus, our finding only *APOE* ε4 indicates that there are few or no other common variants in the genome with an effect on survival as strong as seen in *APOE* region.

We then turned to the UK Biobank data. We applied our method to individuals of British ancestry whose data passed our filters; of these, 88,595 had death information available for their father and 71,783 for their mother. We analyzed 590,437 genotyped autosomal variants, applying similar quality control measures as with the GERA dataset (see Materials and Methods). We tested for significant changes in allele frequencies with father’s age at death and mother’s age at death stratified in eight 5-year interval bins. As in the GERA dataset, no inflation was observed in the quantile-quantile plots (Figure S14).

Consistent with recent studies [33, 34], the variants showing a genome-wide significant change in allele frequency with father’s age at death (*P* < 5×10^−8^) reside within a locus containing the nicotine receptor gene *CHRNA3* (Figure 3A). The A allele of the *CHRNA3* SNP rs1051730 (G, major allele; A, minor allele) has been shown to be associated with increased smoking quantity among individuals who smoke [55]. We observed a linear decrease in the frequency of the A allele of rs1051730 throughout almost all age ranges (Figure 3B). Although it does not reach genome-wide significance, this allele shows a similar trend with age in GERA (*P*~0.0086, Figure S15). We note that 30,819 of the UK Biobank individuals included in the above analysis were genotyped on the UK BiLEVE Axiom array (see Materials and Methods), selected based on lung function and smoking behavior (while the remaining 57,776 samples were genotyped on the UK Biobank Axiom array) [56]. Expectedly, the frequency of the A allele is significantly higher among UK BiLEVE subjects (*P*~2.3×10^−10^), but the age effects are similar across both arrays (*P*~ 0.72).

**Figure 3.**
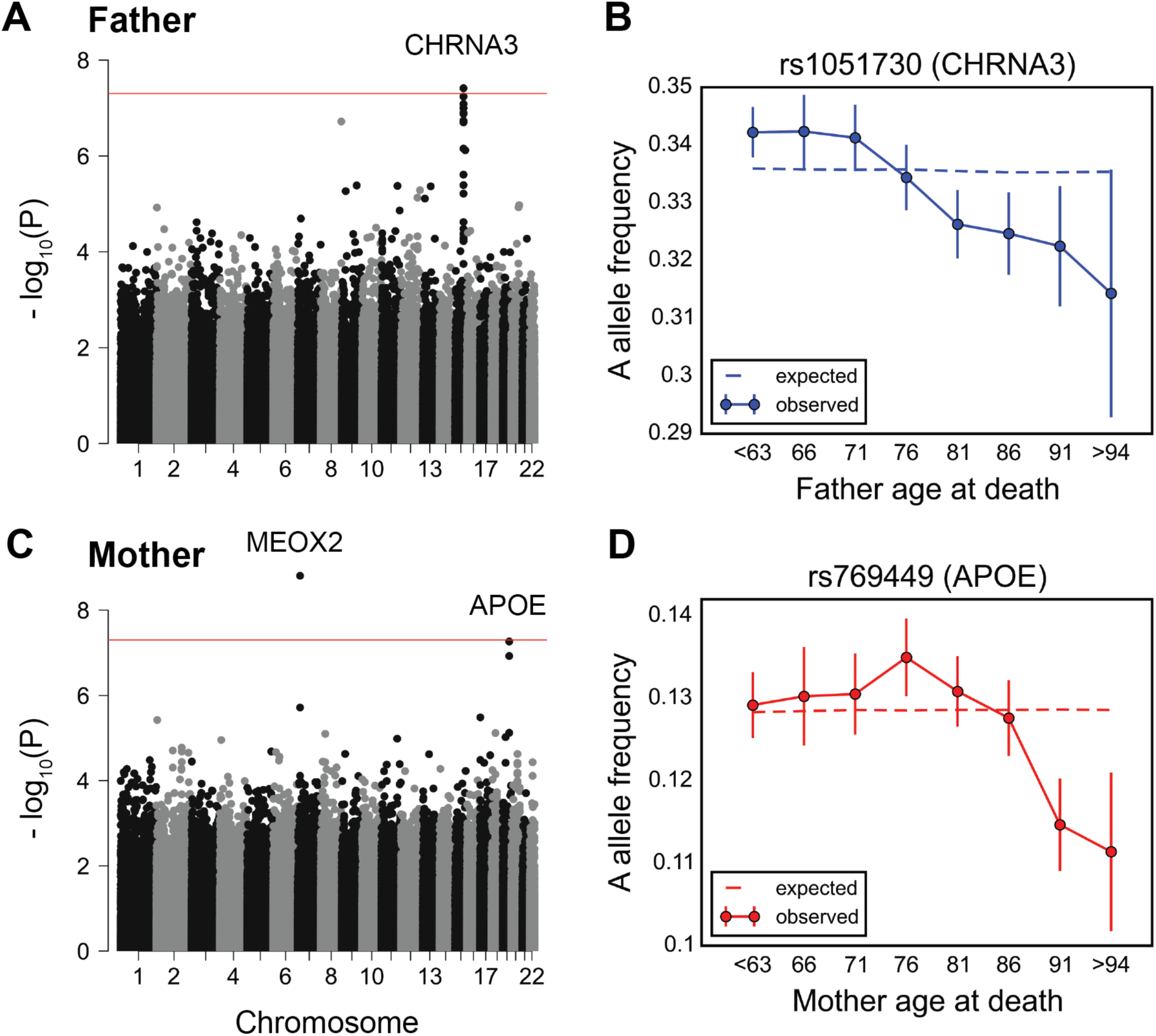
Testing for the influence of single genetic variants on age-specific mortality in the UK Biobank. (A) Manhattan plot of *P* values, obtained from testing for a change in allele frequency with age at death of fathers. (B) Allele frequency trajectory of rs1051730, within *CHRNA3* locus, with father’s age at death. (C) Manhattan plot of *P* values, obtained from testing for a change in allele frequency with age at death of mothers. (D) Allele frequency trajectory of rs769449, within the *APOE* locus, with mother’s age at death. Red lines in (A) and (C) mark the *P* = 5×10^−8^ threshold. Data points in (B) and (D) are mean frequencies of the risk allele within 5- year interval age bins (and 95% confidence interval), with the center of the bin indicated on the x-axis. The dashed line shows the expected frequency based on the null model, accounting for confounding batch effects and changes in ancestry (see Materials and Methods).

For mother’s age at death, a SNP in a locus containing the *MEOX2* gene reached genome-wide significance (Figure 3C). The C allele of rs4721453 (T, major allele; C, minor allele) increases in frequency in the age bin centered at 76 years old (Figure S16), i.e., there is an enrichment among individuals that died at 74 to 78 years of age, which corresponds to a deleterious effect of the C allele in this period. The trend is similar and nominally significant for other genotyped common SNPs in moderate linkage disequilibrium with rs4721453 (Figure S16). Also, the signal for rs4721453 remains nominally significant when using subsets of individuals genotyped on the same genotyping array: 44,552 individuals on the UK Biobank Axiom array (*P*~7×10^−5^) and 25,231 individuals on the UK BiLEVE Axiom array (*P*~10^−4^). These observations suggest that the result is not due to genotyping errors, but it is not reproduced in GERA (*P*~0.023, Figure S17) and so it remains to be replicated. *APOE* variants were among the top nominally significant variants (*P*~10^−7^) (Figure 3C). At the *APOE* SNP rs769449 (G, major allele; A, minor allele), there is an increase in the frequency of A allele at around 70 years old before subsequent decrease (Figure 3D). This pattern is consistent with our finding in GERA (of a monotonic decrease beyond 70 years of age), considering the difference in patterns expected between allele frequency trends with age among survivors versus individuals who died (Figure S3).

We note that by considering parental age at death of the UK Biobank participants–as done also in [33-35]–we introduce a bias towards older participants, who are more likely to have deceased parents (Figure S18). We confirmed that our top signals are not significantly affected after adjusting for age of the participants (among other potential confounders including participants’ sex, birth year and socioeconomic status, as measured by the Townsend deprivation index): results remain similar for the *MEOX2* SNP rs4721453 (*P*~2.1×10^−9^), *APOE* SNP rs769449 (*P*~1.5×10^−6^), and *CHRNA3* SNP rs1051730 (*P*~1.8×10^−6^ and *P*~4.3×10^−9^ treating paternal age at death as a categorical and an ordinal variable, respectively).

We further tested for trends in allele frequency with parental age at death that differ between fathers and mothers, focusing on 62,719 individuals with age at death information for both parents. No variant reached genome-wide significance level (Figure S19A). The rs4721453 near the *MEOX2* gene and *APOE* variant rs769449 show nominally significant sex effects (*P*~7×10^−8^ and *P*~2×10^−3^, respectively), with stronger effects in females (Figure S19B). Variants near the *CHRNA3* locus are nominally significant when using the model with parental age at deaths treated as ordinal variables (rs11858836, *P*~6×10^−4^), with stronger effects in males (Figure 18B).

### Testing for changes in allele frequency at trait-associated variants

We next turned to sets of genetic variants that have been associated with polygenic traits, rather than individual genetic variants. We focused on 42 polygenic traits, including disease risk and traits of evolutionary importance such as puberty timing, for which a large number of common variants have been mapped in GWAS (see Table S1 for the list of traits and number of loci) [37-40]. For each individual and each trait, we calculated a polygenic score based on the genetic variants that reached genome-wide significance level for association, and then tested whether this polygenic score, or its squared difference from the mean in the case of stabilizing selection, is associated with survival (after controlling covariates; see Materials and Methods).

We first applied the Cox proportional hazards model in the UK Biobank for parental lifespan, focusing on the participants whose genetic ancestry is British and who reported their father’s or mother’s age or ages at death (114,122 and 116,323 individuals, respectively). We then compared the results with our approach of testing for changes in the polygenic score across parental ages at death. We further analyzed two data sets for replication purposes: participants of the UK Biobank of non-British ancestry (29,511 and 30,3722 individuals reporting father’s or mother’s age information, respectively) and the GERA cohort.

Using the Cox model, the score for several traits showed significant association with father’s survival after accounting for multiple testing (Figure 4A, Table 1): total cholesterol (TC, *P*~4.3×10^−11^), low-density lipoproteins (LDL, *P*~8.1×10^−9^), body mass index (BMI, *P*~1.8×10^−8^), and coronary artery disease (CAD, *P*~9×10^−6^), consistent with two recent studies [34, 35]. In addition, we uncovered significant association between the polygenic score for puberty timing (*P*~6.2×10^−6^); in this analysis, we use age at menarche associated variants in females, motivated by the high genetic correlation between the timing of puberty in males and females [57]). A higher score for puberty timing was associated with longer paternal survival (per year hazard ratio of 0.96) (Table 1), indicating that variants delaying puberty timing are associated with a higher chance of survival, consistent with epidemiological studies suggesting early puberty timing to be associated with adverse health outcomes [58]. For all other traits, a higher score was negatively associated with paternal survival: one standard deviation (SD) hazard ratio of 1.09 for TC, 1.08 for LDL and 1.22 for BMI (Table 1). With the exception of lipid traits, the effects on survival were not significantly changed after accounting for the effect of the polygenic score of another trait (Figure S20). This is especially relevant to BMI and puberty timing, where there is substantial genetic overlap [38]; the per year hazard ratio was 0.97 for the puberty timing score (*P*~4.8×10^−4^), after adjusting for the BMI score.

**Figure 4.**
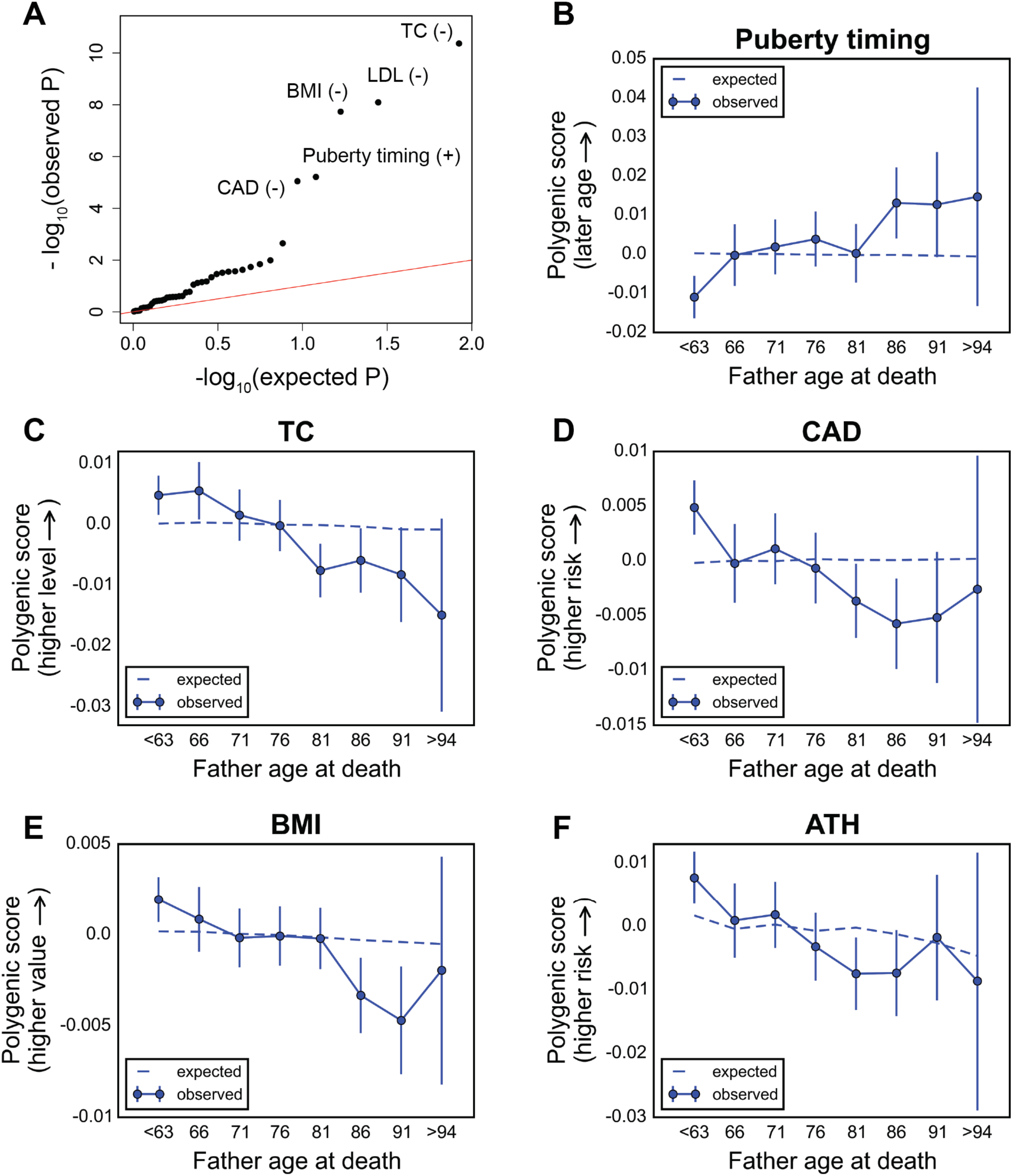
Testing for the influence of set of trait-associated variants on survival of the fathers of UK Biobank participants. (A) Quantile-quantile plot for association between the polygenic score of 42 traits (see Table S1) with father’s survival, using the Cox model. The red line indicates the distribution of the *P* values under the null model. Signs ‘+’ and signs ‘-’ indicate protective and detrimental effects associated with higher values of polygenic scores, respectively. See Table S2 for *P* values and hazard ratios for all traits. (B)-(F) Trajectory of polygenic score with age at death of fathers for top traits associated with paternal survival (only independent signals are shown, see Figure S20): puberty timing (using age at menarche associated variants) in males (B), total cholesterol (C), coronary artery disease (D), body mass index (E), and asthma (F). Data points in (B)-(F) are mean polygenic scores within 5-year interval age bins (and 95% confidence interval), with the center of the bin indicated on the x-axis. The dashed line shows the expected score based on the null model, accounting for confounding batch effects, changes in ancestry, and participant’s age, sex, birth year, and the Townsend index (a measure of socioeconomic status).

**Table 1.** Association with paternal and maternal survival among the UK Biobank participants of British ancestry under the Cox model.

| Trait | Scaling of effect | Father |  |  | Mother |  |  |
| --- | --- | --- | --- | --- | --- | --- | --- |
|  |  | Effect size (s.e.) | HR* | P value | Effect size (s.e.) | HR* | P value |
| Puberty timing | 1 year | -0.0363 (0.0080) | 0.96 | $6.2 \times 10^{-6}$ | -0.0278 (0.0090) | 0.97 | 0.0020 |
| AFB | 1 year | -0.0398 (0.0180) | 0.96 | 0.027 | -0.0639 (0.0200) | 0.94 | 0.0014 |
| ATH | 1 unit log-odds | 0.0279 (0.0109) | 1.03 | 0.010 | 0.0149 (0.0121) | 1.02 | 0.22 |
| BMI | 1 SD | 0.1996 (0.0355) | 1.22 | $1.8 \times 10^{-8}$ | 0.0823 (0.0395) | 1.08 | 0.036 |
| CAD | 1 unit log-odds | 0.0784 (0.0177) | 1.08 | $9.0 \times 10^{-6}$ | 0.0892 (0.0196) | 1.09 | $5.2 \times 10^{-6}$ |
| HDL | 1 SD | -0.0340 (0.0139) | 0.97 | 0.014 | -0.0605 (0.0154) | 0.94 | $8.9 \times 10^{-5}$ |
| LDL | 1 SD | 0.0806 (0.0140) | 1.08 | $8.1 \times 10^{-9}$ | 0.0844 (0.0155) | 1.09 | $5.2 \times 10^{-8}$ |
| TC | 1 SD | 0.0901 (0.0137) | 1.09 | $4.3 \times 10^{-11}$ | 0.0679 (0.0152) | 1.07 | $7.8 \times 10^{-6}$ |

\*Hazard ratio.

Using our approach instead, that is considering the father’s age at death, led to very similar results. Specifically, all traits significantly associated with paternal survival showed a significant change in polygenic score with father’s age at death, using the model with parental age at deaths treated as ordinal variables (Figure S21): TC (*P*~8.7×10^−9^), CAD (*P*~3.3×10^−8^), puberty timing (*P*~1.6×10^−7^), LDL (*P*~8.6×10^−7^) and BMI (*P*~3.4×10^−6^). In addition, we uncovered significant changes in polygenic score with father’s age at death for asthma (ATH, *P*~9.4×10^−5^) and triglycerides (TG, *P*~4.4×10^−4^, the effect of which does not seem to be distinct from other lipid traits, Figure S20). The score for puberty timing increased monotonically with the father’s age at death (Figure 4B), indicative of a protective effect of later predicted puberty timing, whereas all other traits with significant signal showed a monotonic decline in score with age (Figure 4C-F).

For mothers, as in fathers, in a Cox survival model, scores for TC, CAD and LDL were significantly associated with survival, with similar hazard ratios (Figure 5A and Table 1): one SD hazard ratio of 1.09 for LDL (*P*~5.2×10^−8^), 1.09 for CAD (*P*~5.2×10^−6^) and 1.07 for TC (*P*~7.8×10^−6^). In addition, the HDL score was associated with maternal survival (one SD hazard ratio of 0.94, *P*~8.9×10^−5^). Also, suggestive evidence was detected for protective effects of increased predicted age at first birth (AFB) (per year hazard ratio of 0.94, *P*~1.4×10^−3^), as well as predicted puberty timing (per year hazard ratio of 0.97, *P*~1.9×10^−3^) (Figure 5A and Table 1). Other than the LDL and TC, all signals seem to be distinct (Figure S20), including for puberty timing and AFB, despite the genetic correlation between the two phenotypes [39].

**Figure 5.**
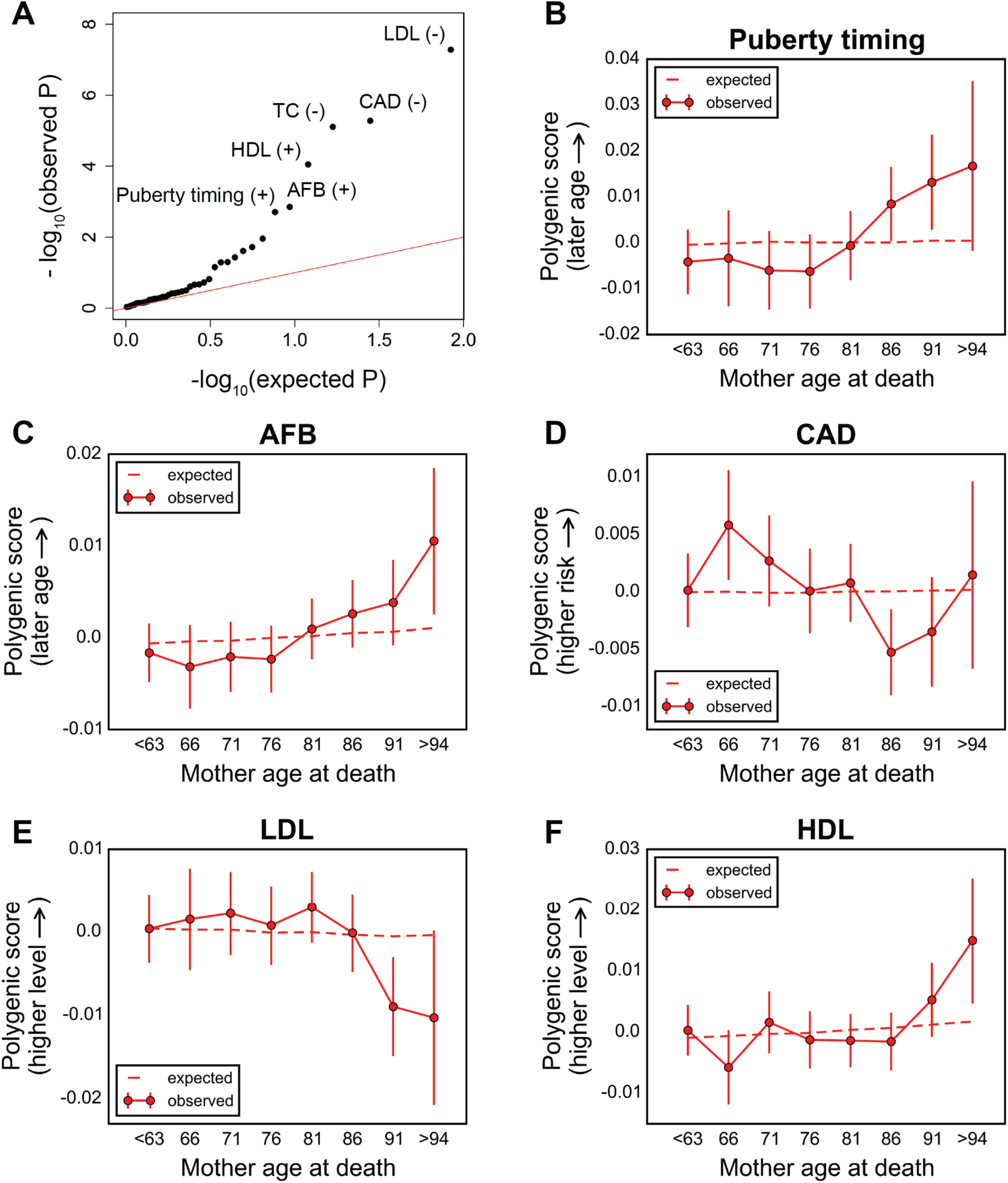
Testing for the influence of set of trait-associated variants on survival of the mothers of UK Biobank participants. (A) Quantile-quantile plot for association between the polygenic score of 42 traits (see Table S1) with mother’s survival, using the Cox model. The red line indicates the distribution of the *P* values under the null. Signs ‘+’ and ‘-’ signs indicate protective and detrimental effects associated with higher values of polygenic scores, respectively. See Table S2 for *P* values and hazard ratios for all traits. (B)-(F) Trajectory of polygenic score with age at death of mothers for top traits associated with maternal survival (only independent signals are shown, see Figure S20): puberty timing (B), age at first birth (C), coronary artery disease (D), low-density lipoproteins (E), and high-density lipoproteins (F). Data points in (B)-(F) are mean polygenic scores within 5-year interval age bins (and 95% confidence interval), with the center of the bin indicated on the x-axis. The dashed line shows the expected score based on the null model, accounting for confounding batch effects, changes in ancestry, and participant’s age, sex, birth year, and the Townsend index (a measure of socioeconomic status).

Applying our approach to maternal age of death instead, puberty timing and AFB were the top signals (*P*~2.2×10^−4^ and *P*~3.1×10^−3^, respectively, Figure S21). Higher polygenic scores for puberty timing were enriched among longer-lived mothers (Figure 5B), as seen for fathers. Similarly, the score for AFB increased with mother’s age at death (Figure 5C), indicating an association between variants that delay AFB and longer lifespan. Scores for CAD, LDL and HDL did not show significant monotonic change across mother’s age at death bins (*P*~7.7× 10^−3^,P~0.058 and *P*~0.35, respectively); however, the trends were suggestive of subtle age dependent effects, with an effect of CAD score in middle-age and late-onset effects of LDL and HDL scores (Figure 5D-F). Testing for age by sex interactions, the TC and CAD score trends with parental age at death were significantly different between fathers and mothers (*P*~4.0×10^−4^ and *P*~7.4×10^−4^, respectively, Figure S22).

To further investigate the age dependency of the effects, we plotted polygenic scores among parents who had survived up to a given age, as compared to the trends with parental ages at death (Figures S23 and S24). All traits associated with paternal survival seemingly show more pronounced effects in middle-ages (Figure S23). Similar patterns were observed for maternal survival associated traits, expect for LDL and HDL with more pronounced late-age effects (Figure S24). We also compared the hazard ratios for ages at death of ≤ 75 and > 75 years (Materials and Methods), similar to a recent study [33]. Consistent with trends in scores with parental age, among the traits associated with paternal survival, almost all traits had stronger effects among younger fathers, particularly for CAD (Table S3): one SD hazard ratio of 1.14 for younger fathers (*P*~2.6×10^−9^), and 0.99 for older fathers (*P*~0.70). Unlike in fathers, in mothers, TC, LDL, and HDL scores had more pronounced late-age effects (Table S3): for TC, one SD hazard ratio of 1.03 for younger mothers (*P*~0.15) and 1.1 for older mothers (*P*~1.4× 10^−6^), and for LDL one SD hazard ratio of 1.05 for younger mothers (*P*~0.03) and 1.12 for older mothers (*P*~3.3×10^−8^).

Next, we sought to replicate the top associations observed among the UK Biobank participants of British ancestry (discovery cohort) in two other data sets: participants of the UK Biobank of non-British ancestry, and the GERA cohort. Applying the Cox model using parental survival for UK Biobank participants of non-British ancestry, the direction of hazard ratios for all traits (as well as the estimated values for most traits) were consistent with the discovery cohort, for both fathers and mothers (Table S4). The congruence of results in two cohorts with different ancestries suggests that our top signals are not false-positives caused by poor control for population structure. In the GERA cohort, we tested whether polygenic scores change with the age of the participant, similar to our approach for individual genetic variants in this cohort. All top signals except AFB had directionally consistent effects with the discovery cohort (Table S5). Of particular interest, the strongest signal was an increase in the polygenic score for puberty timing with age of the participants (*P*~0.0067, Figure S25).

In the discovery cohort, we further investigated if there are significant changes in the squared difference of polygenic scores with parental ages at death, as might be expected if the mean value of the trait leads to the highest change of survival. No trait showed evidence of such stabilizing selection (Figure S26).

## Discussion

We introduced a new approach to identify genetic variants that affect survival to a given age and thus to directly observe viability selection ongoing in humans. Attractive features of the approach include that we do not need to make a decision a priori about which loci or traits matter to viability and focus not on an endpoint (e.g., survival to an old age) but on any shift in allele frequencies with age, thereby learning about the ages at which effects are manifest and possible differences between sexes.

To illustrate the potential of our approach, we performed a scan for genetic variants that impact age-specific mortality in the GERA and the UK Biobank cohorts. We only found a few individual genetic variants, the majority of which were identified in previous studies. This result is in some ways expected: available data only provide high power to detect effects of common variants (>15-20%) on survival (Figure 1), yet if these variants were under viability selection, we would not expect them to be common, short of strong balancing selection due to trade-offs between sexes, ages or environments. As sample sizes increase, however, the approach introduced here should provide a comprehensive picture of viability selection in humans. To illustrate this point, we repeated our power simulation with 500,000 samples, and found that we should have high power to detect the trends for alleles at a couple percent frequency (Figure S27).

Already, however, this application raises a number of interesting questions about the nature of viability selection in humans. Notably, we discovered only a few individual variants influencing viability in the two cohorts, most of which exert their effect late in life. On first thought, this finding may suggest such variants to be neutrally-evolving. We would argue that if anything, our findings of only a few common variants with large effects on survival late in life suggest the opposite: that even variants with late onset effects have been weeded out by purifying selection. Indeed, unless the number of loci in the genome that could give rise to such variants(i.e., the mutational target size) is tiny, other variants such as *APOE* ε4 must often arise. That they are not observed when we have very high power to detect them suggests they are kept at lower frequency by purifying selection. Why might they be selected despite affecting survival only at old ages? Possible explanations include that they decrease the direct fitness of males sufficiently to be effectively selected (notably given the large, recent effective population size of humans [59]) or that they impact the inclusive fitness of males or females. If this explanation is correct, it raises the question of why *APOE* ε4 has not been weeded out. We speculate that the environment today has changed in such a way that has made this allele more deleterious recently. For example, it has been proposed that the evolution of this allele has been influenced by changes in physical activity [60] and parasite burden [61].

Considering 42 traits that have been investigated by GWAS, we found a number of cases in which the mean polygenic score changes with age. Of course, detecting an effect of age on the traits does not imply that these are the phenotypes under viability selection, as the variants that contribute likely have pleiotropic effects on other traits [37]. Nonetheless, it is perhaps not surprising that we found detrimental effects of higher genetically predicted TC, LDL, BMI and the risk of CAD on survival, as these phenotypes are studied in GWAS precisely because of their adverse health effects. Intriguingly, however, we also found associations for fertility traits, notably protective effects of later predicted puberty timing and AFB. If these findings reflect life-history trade-offs (e.g., longer lifespan at the cost of delayed reproduction), they may help to explain the persistence of extensive variation in such fitness-correlated traits [62, 63]. Intriguingly, we do see a negative correlation between genetically predicted AFB and number of siblings of the UK Biobank participants, a proxy for the fertility of their parents (*P*~4.2×10^−8^, Figure S28), consistent with previous reports of a genetic correlation between AFB and the number of children ever born [21, 39]. These findings underscore that consideration of survival or fertility effects alone does not allow one to infer whether the net effect of a variant or set of variants is beneficial. Instead, to convert effects on viability, such as those detected here, or effects on fertility reported elsewhere [22, 23] into an understanding of how natural selection acts on an allele requires a characterization of its effects on all components of fitness (including potentially inclusive fitness).

In this regard, it is also worth noting that while our method is designed to detect changes in allele frequencies (and in polygenic scores) caused by genetic effects on age-specific survival, such changes could in principle also arise from effects on other components of fitness. For example, if the frequency of a genetic variant in a population decreases over decades due to an effect on fertility, its frequency would increase with the age of surviving individuals sampled at a given time (as in the GERA cohort). This confounding is less of an issue when considering effects on the age at death (what we measured in the UK Biobank). Nonetheless, even in the UK Biobank, fertility effects may manifest as effects on age of death; for example, because when sampling a cohort of children, parents with later ages at death are possibly born earlier (Figure S29). To this end, in the UK Biobank, we account for changes in allele frequencies with year of birth of the participants themselves (ideally we would want to condition on parents born at similar times, which we cannot do; instead, we used birth year of the participants as an estimator for birth year of the parents). Thus, we believe our results in the UK Biobank not to be confounded by fertility effects. Moreover, a number of our findings in this study are consistent with prior knowledge of effects on survival, such as those for disease risk variants like *APOE* ε4. Nonetheless, some caution is required in interpreting trends with age as strictly reflecting viability effects.

Also of interest are the marked differences between males and females in our analysis of mothers and fathers of individuals in the UK Biobank. The differences between sexes are most notable at the *CHRNA3* locus, which shows a strong effect only in fathers, and sets of genetic variants associated with risk of coronary artery disease and cholesterol levels, which exhibit different age-dependent effects between fathers and mothers. Results for the *CHRNA3* locus, in which variants are associated with the amount of smoking among smokers, may reflect a gene-by-environment interaction rather than a sex effect. Consistent with a more pronounced effect on male than female age at death, smoking prevalence in men has been consistently higher than women over the past few decades in the UK: from 1970 to 2000, smoking prevalence decreased from around 70% to 36% in middle-aged men, compared to from around 50% to 28% in middle-aged women [64].

Many of these questions can soon be addressed, by applying approaches such as ours to the millions of samples in the pipeline (such as the UK Biobank [65], the Precision Medicine Initiative Cohort Program [66], and the Vanderbilt University biobank (BioVU)[67]), in which the viability effects of rare as well as common alleles can be examined. These analyses will provide a comprehensive answer to the question of which loci affect survival, helping to address long-standing open questions such as the relative importance of viability selection in shaping genetic variation and the extent to which genetic variation is maintained by fitness trade-offs between sexes or across ages.

## Materials and Methods

### 1. Datasets

#### 1.1. GERA cohort

We performed our analyses on the data for 62,318 participants of the Kaiser Permanente Northern California multi-ethnic Genetic Epidemiology Research on Adult Health and Aging (GERA) cohort, self-reported to be “White-European American”, “South Asian”, “Middle-Eastern” or “Ashkenazi” but no other ethnicities, among a list of 23 choices on the GERA survey, and genotyped on a custom array at 670,176 SNPs designed for Non-Hispanic White individuals [31, 32]. We determined the age of the participants and the number of years they were enrolled in the Kaiser Permanente Medical Insurance Plan at the time of the survey (year 2007).

#### 1.2. UK Biobank

We performed our analyses on the data for 152,729 participants of the UK Biobank study, focusing on 120,286 individuals identified to be “British” by genetic analysis, and all other individuals for replication. They were genotyped on the UK Biobank Axiom or the UK BiLEVE Axiom SNP arrays at a total of 847,441 SNPs in the interim release [56, 65].

### 2. Quality control (QC)

#### 2.1. GERA cohort

We used PLINK v1.9 [68] to remove individuals with missing sex information or with a mismatch between genotype data and sex information, individuals with <96% call rate, and related individuals. We validated self-reported European ancestries using principal component analysis (PCA), see below, and removed individuals identified as non-European (Figures S4 and S5). In the end, 57,696 individuals remained.

Using PLINK, we removed SNPs with <1% minor allele frequency, SNPs with <95% call rate, and SNPs failing a Hardy-Weinberg equilibrium test with *P* < 10^−8^ (filtering based on HWE test could potentially exclude true signals of viability selection, if selection coefficients were very large [69], but this possibility is much less likely than genotyping error). We additionally tested for a correlation between age (or sex) and missingness, which can induce artificial change in the allele frequencies as a function of age (or sex). We thus removed SNPs showing a significant age-missingness or sex-missingness correlation, defined as a chi-squared test with *P* < 10^−7^. After these steps, 599,659 SNPs remained.

We imputed the genotypes of the filtered GERA individuals using post-QC SNPs, and using the 1000 Genomes phase 3 haplotypes as a reference panel [70]. We phased observed genotypes using EAGLE v1.0 software [71]. The inferred haplotypes were then passed to IMPUTE2 v2.3.2 software for imputation in chunks of 1Mb, using the default parameters of the software [72]. To gain computational speed, variants with <0.5% minor allele frequency in the 1000 Genomes European populations were removed from the reference panel. This step should not affect our analysis because our statistical model is not well powered for rare variants, given the GERA data sample size. We called imputed genotypes with posterior probability >0.9, and then filtered the imputed genotypes, removing variants with IMPUTE2 info score <0.5 and with minor allele frequency <1%. We also used imputation with leave-one-out approach [73] to impose a second stage of QC on genotyped SNPs, removing SNPs that were imputed back with high reported certainty (info score >0.5) and with <90% concordance between the imputed and the original genotypes. These yielded a total of 9,010,280 imputed and genotyped biallelic SNPs and indels.

For our analysis of the *APOE* alleles (ε2, ε3 and ε4) which are defined by rs7412 and rs429358 SNPs [54], given the lack of tag SNPs for all three alleles, we kept a subset of 38,703 individuals with no poorly-imputed genotypes for these two SNPs, for whom the count of each *APOE* allele could be determined unambiguously.

#### 2.2. UK Biobank

In the UK Biobank, we obtained sets of genotype calls and the output of imputation as performed by the UK Biobank researchers [56, 74]. We first applied QC metrics to the autosomal genotyped SNPs, focusing on the individuals of British genetic ancestry. We used PLINK to remove SNPs with <1% minor allele frequency, SNPs with <95% call rate, and SNPs failing a Hardy-Weinberg equilibrium test with *P* < 10^−8^. These filters were applied separately to SNPs genotyped on the UK Biobank Axiom and the UK BiLEVE Axiom arrays. Then, we divided the genotyped SNPs into three sets (SNPs specific to either array and shared SNPs) and then performed additional QC on each set separately: we removed SNPs with significant allele frequency difference between genotyped and imputed calls (chi-squared test *P* < 10^−5^) and SNPs showing a significant correlation between missingness and age or sex of the participants, as well as with participants’ father’s or mother’s age at death (chi-squared test *P* < 10^−7^). We then extracted this list of SNPs from the imputed genotype files available from the UK Biobank (we did not use the full set of imputed genotypes). From this set, we removed SNPs with <1% minor allele frequency, SNPs with <95% call rate, and SNPs failing a Hardy-Weinberg equilibrium test with *P* < 10^−8^, yielding 590,437 SNPs. For variants influencing quantitative traits, we first extracted them from imputed genotypes, and then imposed the same QC measures as above. For individuals of non-British ancestry, we first extracted the trait influencing variants from imputed genotypes, and then removed SNPs with <1% minor allele frequency and SNPs with <90% call rate.

Each participant was asked to provide the survival status and age of their father and their mother on each assessment visit. For each participant that reported an age at death of father and/or mother, we averaged over the ages reported at recruitment and any subsequent repeat assessment visits, and used PLINK to exclude individuals with >5 year variation in their answers across visits (around 800 individuals). For those reporting their parents to be alive, the latest assessment visit was considered. We also removed adopted individuals, individuals with a mismatch between genotype data and sex information, and individuals with missing values for the covariates, resulting in 88,595 individuals of British ancestry with age at death information for their father, 71,783 individuals for their mother, and 62,719 individuals for both parents. For the survival analyses, we further removed individuals with evidently invalid survival status, particularly parental ages at death values smaller than their age when still alive, resulting in 114,122 and 116,323 individuals of British ancestry with paternal and maternal survival information, respectively. With similar quality control measures, 29,511 and 30,372 individuals with non-British ancestry with paternal and maternal survival information, respectively, were analyzed.

### 3. Principal Component Analysis

We performed PCA, using the EIGENSOFT v6.0.1 package with the fastpca algorithm [75, 76], for two purposes: (i) as a quality control on individuals to validate self-reported European ancestries (only in GERA dataset), and (ii) to correct for population structure in our statistical model (for individuals in the UK Biobank of non-British ancestry, we used the PCs provided with the data).

#### 3.1. European ancestry validation

We used more stringent QC criteria specifically for the PCA, compared to the QC steps described above. We filtered a subset of 157,277 SNPs in GERA, retaining SNPs shared between the datasets and the 1000 Genomes phase 3 data, removing non-autosomal SNPs, SNPs with <1% minor allele frequency, SNPs with <99% call rate, and SNPs failing a Hardy-Weinberg equilibrium test with *P* < 10^−6^. We then performed LD-pruning using PLINK with pairwise *r*^2^ <0.2 in windows of 50 SNPs shifting every 10 SNP. We used these SNPs to infer principal components for the 1000 Genomes phase 3 data [70]. We then projected individuals onto these PCs. We observed that the majority of individuals have European ancestry, and marked individuals with PCs deviating from the population mean, for any of the first six PCs, as nonEuropean (Figures S4 and S5).

#### 3.2. Control for population structure

After the main QC stage, additional QC steps (as in section 3.1) were implemented for PCA. In the UK Biobank, we also removed inversion variants on chromosome 8 which otherwise dominate the PC2 (not shown). A subset of 156,721 SNPs in GERA and 207,657 SNPs in the UK Biobank was then used to infer PCs for individuals passing QC (Figure S1). The first 10 PCs were used as covariates in our statistical model.

### 4. Quantitative Traits

We downloaded the list of variants contributing to 39 traits (all traits but age at menarche, age at first birth and age at natural menopause) and their effect sizes recently described in Pickrell et al. [37], from: https://github.com/PickrellLab/gwas-pw-paper/tree/master/all_single. For age at menarche, we used the variants and effect sizes recently identified by Day et. al [38]. We used variants associated with age at first birth from Barban et al., identified in either sex-specific analyses or analyses of both sexes and used the effect sizes estimated in the combined analysis [39]. We used age at natural menopause associated variants and their effect sizes from Day et al. [40]. For all traits, we used variants that were genotyped/imputed with high quality in our data (see Table S1).

### 5. Statistical Model

#### 5.1. An individual variant

Using a logistic regression we predict the genotype of individual *j* (the counts of an arbitrarily selected reference allele, *G_ij_* = 0, 1 or 2) at variant *i*, using the individual’s ancestry, the batch at which the individual was genotyped, and individual’s age (as well as sex, see below) as explanatory variables. Specifically, the distribution of *G_ij_* is *Bin*(2, *p_ij_*), where *p_ij_*, the probability of observing the reference allele for individual *j* at variant *i*, is related to explanatory variables as:

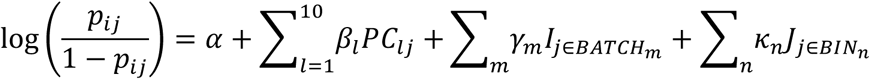

where *β_l_* is the effect of principal component *l* (to account for population structure),*γ_m_* is the effect of being in batch *m* (to account for potential systematic differences between genotyping packages), *K_n_* is the effect of being in age bin *n*, obtained by regression across individuals with non-missing genotypes at variant *i*, and *I* and *J* are indicator variables for the genotyping batch and age bin, respectively. In the version of the model in which we treat age as an ordinal variable, we replace *J* age bin variables with one age variable. In the GERA dataset, age binning is over the age of the participants in 14 categories, from age 19 onwards, in 5-year intervals. For replication purposes, we further binned the ages in 7 categories, in 10-year intervals, to boost our power by increasing the sample size per bin, particularly for younger age bins. In the UK Biobank, we binned the age at death of father or mother over 8 categories, from age 63 onwards, in 5-year intervals. In the UK Biobank, we included all ages at death below 63 in one age bin to minimize the potential noise caused by accidental deaths at young ages.

We tested for an effect of age categories by a likelihood ratio test with a null model using only the covariates (PCs and batch terms) (*H*_0_: *K_n_* = 0, for all *n*) and an alternative also including age terms as predictors (*H*_1_: *κ_n_* ≠ 0, for at least one *n*):

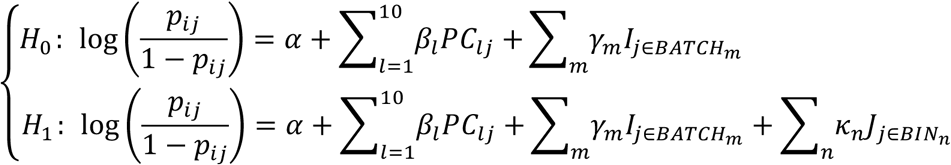

To test for age by sex effects in GERA we included two sets of additional predictors. The first consists in two indicator variables for sex, *K*_male_ and *K*_female_, which are included to capture possible sex effects induced by potential genotyping errors or missmapping of sex chromosome linked alleles (we note that because of Hardy-Weinberg equilibrium, mean allele frequency difference between males and females are not expected). The second set of predictors consists in age by sex terms, *J*×*K*. We then compare a model with age and sex terms as predictors to a model also including age by sex terms. To test for sex effects in the UK Biobank, we compared a model with both father and mother age terms separately as predictors to a model with one set of age categories for average age at death of both parents, only for individuals reporting the age at death for both parents. In all models PCs and batch terms were incorporated as covariates. For the top SNPs in the UK Biobank, we additionally tested models also including as covariates the participants’ age, sex, birth year, and the Townsend index (a measure of socioeconomic status). For rs1051730, we also tested whether allele frequencies or trends in allele frequencies with the father’s age at death vary significantly across the UK Biobank genotyping arrays after adjusting for population structures, using similar models as described above.

#### 5.2. Set of variants

As for the model described above for an individual variant, we investigated age and age by sex effects on quantitative traits for which large number of large common genetic variants have been identified in genome-wide association studies (GWAS). For a given trait, we used a linear regression with the same covariates and predictors as for the model for an individual variant, to predict the polygenic score for individual *j, S_j_*, by summing the previously estimated effect of single variants assuming additivity and that the effect sizes are similar in the GWAS panels and the cohorts considered here:

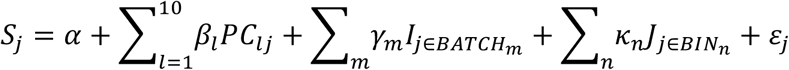

*S_j_* is calculated as ∑*a_i_G_ij_* + ∑2*a_i_q_i_* (standardized to mean 0), where the first sum is across variants with non-missing genotypes, *a_i_* is the effect size for the arbitrary selected reference allele at variant *i*, and the second sum is across the variants with missing genotypes estimating their contribution assuming Hardy-Weinberg equilibrium where *q_i_* is the frequency of the alternate allele. Likelihood ratio tests, as described above, were used to test for age and age by sex effects. In the UK Biobank, we additionally adjusted for participants’ age, sex, birth year, and the Townsend index.

To evaluate the possibility of stabilizing selection on a trait, we applied the same model, but instead of the polygenic score, regressed the squared difference of the score from the mean in each bin, 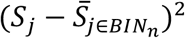, on the predictors, where 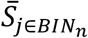 is the mean score in the age bin to which individual *j* belongs.

##### Survival analysis

We also used the Cox proportional hazards model [45] to evaluate the association between polygenic scores and parental survival in the UK Biobank. Compared to the model described above, this approach presents the advantage of allowing data from participants with alive parents to be incorporated, but has the disadvantage of assuming fixed effects across all ages. Under this model, at a given time *t* (age in our application):

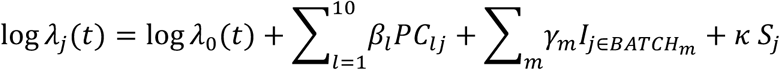

where *λ* is the hazard rate (probability of death within *t* + *dt* conditional on survival to time *t*) given the covariates, and *λ*_0_ is the baseline hazard rate that describes the risk for individuals with the value of 0 for all predictors. Not shown in the equation above are covariates to adjust for participants’ age, sex, birth year, and the Townsend index. Using the R package ‘Survival’ [77], for a given trait, we tested for a significant effect of polygenic score (*κ* ≠ 0). In addition, to assess the interdependence of detected effects (Figure S20), for each pair of traits [*a, b*], we tested for the effect of the polygenic score for trait *a*, but also incorporated the polygenic score for trait *b* as a covariate in the null model (in addition to the covariates mentioned above).

We further investigated the age-dependency of the effects in the framework of the survival analysis by comparing hazard ratios in two age categories: ages at death of ≤ 75 and > 75 years. For the category of ages at death ≤ 75 years, all parental ages were included in the analysis, and parents with ages at death beyond 75 years were marked as alive. For the category of ages at death > 75 years, only parents who survived beyond 75 years were considered.

All Manhattan and quantile-quantile plots were generated using qqman [78] and GWASTools [79] packages.

### 6. Power simulations

We ran simulations to determine the power of our statistical model to detect deviation of allele frequency trends with age across 14 age categories mimicking the GERA individuals (57,696 individuals with age distribution as in Figure S2) from a null model, which for simplicity was no change in frequency with age, i.e., no changes as a result of age-dependent variation in population structure and batch effects. For a given trend in frequency of an allele with age, we generated 1000 simulated trends where the distribution of the number of the alleles in age bin *i* is Bin(2*N_i_*,*f_i_*), where *N_i_* and *f_i_* are the sample size and the sample allele frequency in bin *i*. We then estimated the power to detect the trend as the fraction of cases in which *P* < 5× 10^−8^, by a chi-squared test.

### 7. Survival simulations

We ran simulations to investigate the relationship between allele frequency with age of the survived individuals and the age of the individuals who died in a cohort. We simulated 2×10^6^ individuals going forward in time in 1 year increments. For each time step forward, we tuned the chance of survival of the individuals based on their count of a risk allele for a given variant such that the number of individuals dying in the increment complies with: (i) a normal distribution of ages at death with mean of 70 years and standard deviation of 13 years, roughly as is observed for parental age at deaths in the UK Biobank, and (ii) a given frequency of the risk allele among those who survive. Specifically, we modeled the survival rate of the population, *S*, as the weighted mean for 2 alleles carriers, *S*_2_, 1 allele carriers, *S*_1_, and non-carriers, *S*_0_:

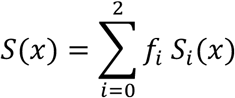

where *f* denotes the frequency of genotypes in the population and *x* denotes the age. *S_i_* and *S* are related: *S_i_*(*x*) = *S*(*x*) *f_i_*(*x*)/*f_i_*, where *f_i_*(*x*) is the genotype frequency among individuals survived up to age x. Given a trend in allele frequency with age, we calculated genotype frequencies with age assuming Hardy-Weinberg equilibrium, and then estimated genotype dependent chance of survival, *S_i_* (*x*), taking *S*(*x*) as the survival function for *N*(70,13^2^).

## Acknowledgments

We thank Guy Sella and members of the Pickrell, Przeworski and Sella labs for helpful discussions and Graham Coop and Jonathan Pritchard for comments on an earlier version of the manuscript. This research has used the UK Biobank Resource (application number 11138), and was funded in part by Columbia University (a Research Initiative in Science and Engineering grant to MP and JKP) and the National Institutes of Health (grant R01MH106842 to JKP and R01GM115889 to Guy Sella). FRD and JRBP and supported by the Medical Research Council [Unit Programme number: MC_UU_12015/2]. These data analyses were approved by the Columbia University Institutional Review Board, protocols AAAQ2700 and AAAP0478.

